# Motor and non-motor sequence prediction is equally affected in children with Developmental Coordination Disorder

**DOI:** 10.1101/2020.04.20.050864

**Authors:** Bertram Opitz, Daniel Brady, Hayley C. Leonard

**Author notes:** Corresponding author (BO). Author Contributions BO contributed to funding acquisition, conceptualization of the study, project administration and supervision, data curation, and writing of the paper and visualisation DB programmed the tasks, acquired and analysed the data, and contributed to writing of the paper and visualisation HCL contributed to funding acquisition, conceptualization of the study, project administration and supervision, and writing of the paper.

## Abstract

Children with Developmental Coordination Disorder (DCD) are diagnosed based on motor difficulties. However, they also exhibit difficulties in several other cognitive domains, including visuospatial processing, executive functioning and attention. One account of the difficulties seen in DCD proposes an impairment in internal forward modelling, i.e., the ability to (i) detect regularities of a repetitive perceptual or motor pattern, (ii) predict future outcomes of motor actions, and (iii) adapt behaviour accordingly. Using electroencephalographic recordings, the present study aimed to delineate these different aspects of internal forward modelling across several domains. To this end, 24 children with DCD and 23 typically-developing children (aged 7-10 years) completed a serial prediction task in the visual, temporal, spatial and motor domains. This task required them to learn short sequences and to indicate whether a sequence was disrupted towards its end. Analyses revealed that, across all domains, children with DCD showed poorer discrimination between intact and disrupted sequences, accompanied by a delayed late parietal positivity elicited by disrupted sequences. These results indicate an impairment in explicit sequence discrimination in DCD across motor and cognitive domains. However, there is no evidence for an impairment in implicit performance on the motor task in DCD. These results suggest an impairment of the updating of an internal forward model in DCD resulting in a blurred representation of that model and consequently in a reduced ability to detect regularities in the environment (e.g., sequences). Such a detailed understanding of internal forward modelling in DCD could help to explain the wide range of co-occurring difficulties experienced by those with a diagnosis of DCD.

## Introduction

Developmental Coordination Disorder (DCD) affects 5-6% of children aged 5-11 and is diagnosed based on motor impairments that significantly impact daily life [1]. Although motor impairments are central to the diagnosis, DCD is characterised by difficulties in several other cognitive domains, including visuospatial processing, executive functioning and attention [2,3]. This leads to a range of adverse individual and societal consequences associated with DCD in terms of poorer physical health [4,5], mental health, [6,7]and reduced academic/employment opportunities and success [8]. The causes of these impairments are poorly understood and, therefore, further research is warranted to bolster successful treatment strategies.

### Internal forward modelling account

One potential common mechanism underlying the difficulties presented in DCD might be a deficit in effectively sequencing information. Sequencing implies the ability to detect regularities or repetitive patterns in an information stream and is also important for predicting future events [9,10]. The ability to correctly predict the end of a sequence of movements is vital for fluent and successful execution of motor actions [11,12]. Prediction enables the reduction of processing resources and preparing the relevant cortices to respond, which in turn improves speed and accuracy of performance [13]. In motor control, it is generally agreed that the process of prediction occurs through the generation of an internal forward model, which uses sensory information from the environment to predict the outcome of movements and allow rapid corrections to these movements if required (e.g. in a situation where an individual’s hand is moving to catch a ball, but the ball bounces and quickly changes trajectory; [11]). More recently, however, an increasing amount of evidence has been reported to suggest that forward internal models are required not only for predicting motor actions, but also for non-motor events (albeit with a reduced level of description, given that they do not require the same precision as motor actions; [9]). Specifically, it has been proposed that a core sensorimotor network, comprising the premotor cortex and connecting parietal lobules as well as the cerebellum, are activated during internal forward modelling across motor, perceptual and cognitive domains [14]. Thus, assuming a deficit in internal forward modelling in DCD, as suggested by a range of research across visuospatial attention and motor control [15] and in both timing prediction [16] and motor and non-motor planning [2,17], could help to explain why difficulties in DCD are seen across different domains. Further supporting evidence for this assumption comes from recent research demonstrating similar deficits in sequencing information across motor and perceptual domains in patients with lesions to premotor, parietal and cerebellar regions (e.g. [9,18,19]). In addition, there is evidence of underactivation of parietal and prefrontal areas in DCD overlapping with brain areas implicated in internal modelling (see [2] for a review). This suggests that deficits in DCD affecting the internal modelling system may be domain-general rather than specific to a certain effector system [15]. The current study therefore aimed to assess whether difficulties across perceptual and motor domains in DCD were associated with atypical neural responses to sequence disruption. This would provide further insight into suggested deficits in internal forward modelling in DCD [15] and into the hypothesised unified sensorimotor network underlying event prediction [14].

Producing an internal forward model requires several stages: (i) the detection or learning of an action sequence, (ii) the ability to use the regularities of the repetitive pattern to be able to predict its outcome and detect any violations, and (iii) the ability to adapt behaviour accordingly [19]. Research into motor control in DCD has shown increased variability or reduced ability to adapt to perturbations in the environment which disrupt action sequences (see [15] for a review). It has been argued that children with DCD appear to have a reduced ability to form internal models for action and to use these stored estimates in a predictive manner. In contrast, according to the internal modelling deficit (IMD) hypothesis, children with DCD have a reduced ability to utilize predictive motor control [3,20]. Consequently, there seems to be a lack of insight into which stage of internal forward modelling is affected. The current study capitalises on the high level of temporal resolution of event-related potentials (ERPs) to understand the neural correlates of different stages of internal modelling, and thus provide a better insight into any disruption in DCD.

### Procedural Learning in DCD

To assess motor sequence detection in DCD, research has mainly focused on procedural learning of motor sequences, or “the ability to combine isolated movements into a single smooth and coherent action” ([21], p.153). Previous studies [3,22,23] have measured procedural learning using the Serial Reaction Time Task (SRTT; [24]), in which participants press buttons corresponding to a stimulus presented in one of four spatial locations, either in a particular sequence (which is not made explicit) or in a random order. In these tasks, reaction times (RTs) to the stimulus should reduce over a number of trials in the sequence condition but not in the random condition if sequence learning is evident. Other studies have used a finger tapping task [25,26], where participants are instructed to memorise and perform a sequence of finger movements repeatedly. Sequence learning is indexed by increasing accuracy over the course of the different blocks. The majority of these studies have reported similar sequence learning between children with DCD and typically-developing controls [23,27] or between DCD and those with other neurodevelopmental disorders [25,26], despite a general slowing of responses in DCD. On the other hand, Gheysen and colleagues [22] found that children with DCD improved their RTs over the course of the training but, unlike a typically-developing control group, did not demonstrate any differences between the sequence and random conditions. This suggests that despite a spared ability to improve motor performance, children with DCD were unable to learn the sequence. These differences between studies may be due to distinct task demands or characteristics of the samples and are, thus, not necessarily indicative of spared or impaired sequence learning in general. Of particular note, however, is the fact that even similar behavioural performance between children with DCD and controls can be subserved by atypical neural functioning. For example, in the study by Biotteau et al. [26], children were tested on a learned finger tapping sequence, which should result in a change in the neural networks recruited for the learned task compared to a novel sequence task. However, children with DCD did not show this difference in neural activity underlying the two sequence tasks, which the authors suggest highlights inefficient or suboptimal processing during sequence learning despite a similar behavioural outcome. Such discrepancies between behavioural and neural functioning thus serve to emphasise the importance of integrating both measures into research into DCD, which was a particular aim of the current study.

Within the internal forward modelling account of DCD, the above findings, i.e., speeding up response times but not differentiating between learned and random sequences, could be explained in two ways: either (i) by impaired sequence learning and/or prediction or (ii) by impaired motor realisation of the predicted sequence. The latter implies that children with DCD are able to correctly predict the next element of a sequence, but they are unable to adequately use this prediction to speed up their motor response. The SSRT cannot easily assess which stage of the internal forward model is most affected in DCD. However, the use of serial prediction tasks (SPTs) allows the different stages to be investigated. SPTs provide regular sequences of pairs of alternating targets which share one stimulus property (e.g. shape) to which participants must attend. In some trials, the sequence is disrupted by repeating the same stimulus twice instead of alternating it with the other stimulus in the pair [9]. Neural responses to these tasks are similar across different perceptual domains [9,14] and the ability to explicitly identify that a sequence violation has occurred is affected by lesions in key areas of the proposed sensorimotor forward modelling network [18]. Applying these tasks to DCD would therefore bring together different bodies of literature investigating internal forward models in both motor and non-motor contexts across populations with typical and atypical functioning of the sensorimotor network. It is to this end that the current study was conducted.

### The current study

The current study was part of a wider research project which also investigated sequencing in linguistic tasks, which is outside the scope of this paper. For the perceptual and motor tasks, the three perceptual SPTs used by Schubotz and colleagues [18] were adopted and integrated into a sequence detection game for children. Both behavioural and neural responses to sequence violations were recorded, using a forced-choice behavioural response at the end of each sequence (i.e. participants indicated whether the sequence had or had not been disrupted). EEG was recorded to assess the P3 response. The P3 is a positive deflection in the ERP occurring around 600ms after the sequence was disrupted at centro-parietal recording sites [28]. It has been shown to be an important signature of stimulus evaluation processes and the updating of a mental model of the environment [29]. Thus, reduced accuracy and an attenuated P3 response to sequence disruptions across all perceptual SPT tasks would be indicative of impaired sequence learning and prediction rather than a selective motor impairment.

The same format was used for a newly-developed visuomotor SPT. In this task, participants pressed buttons on a button box as they lit up in an alternating pattern. Given the correspondence between this task and the perceptual SPTs, a similar pattern of results was expected in response to sequence disruptions in terms of reduced accuracy and an attenuated P3 response to the violation in DCD. The use of ERP to assess sensitivity to sequence disruption in DCD also allows us to identify any potential differences in processing at the neural level, which may not be evident at the motor behaviour level (as in [26]). However, further investigation of the motor SPT was also necessary to help understand mixed results in motor sequence learning research. As in previous studies using the SRTT [22,23,27], the motor SPT included both a perceptual element (i.e. identifying which buttons should be pressed) and a motor execution element (i.e. pressing the buttons). In order to better understand performance on these different elements of the visuomotor SPT, behavioural performance was assessed through both error rates for detecting the sequence disruption and RTs for the production of the motor sequence. This allowed four potential patterns to be explored, each providing a different interpretation of sequence prediction abilities in DCD:

1. The sequence is being learned and executed correctly. This would be revealed through a high accuracy in the identification of sequence violations and a reduction in RTs over the course of the repeated trials within a block of the motor task.
2. There is a general problem with visual sequence perception but not motor sequence production (motor execution). This would be indicated by a reduction in RTs over the course of the repeated trials within a block of the motor task and the inability to explicitly identify the sequence violation in any task.
3. There is a problem with motor sequence production, but not visual sequence perception. This would be demonstrated through a lack of reduction in RTs over the course of the repeated trials within a block of the motor task but a spared explicit identification of sequence violation.
4. There is a problem with both visual sequence perception and motor sequence production as indicated by a lack of reduction in RTs over the course of the repeated trials within a block, as well as an inability to explicitly identify the sequence violation.

The second and fourth scenarios would support the case for a more domain-general sequence prediction deficit in DCD, which would be bolstered by evidence of problems across the other tasks in the battery. Based on the previous mixed evidence, no specific predictions were made about which of these scenarios was more likely within the DCD group.

## Materials and methods

### Participant recruitment and screening

Children between 7-10 years old were recruited into two groups: those with DCD, and typically-developing (TD) controls. Children with DCD were recruited through an advertisement placed on the Facebook pages of a charitable organisation, the Dyspraxia Foundation, and of the researchers’ lab group. The advertisement was also placed on the lab group website and Twitter page. Parents who were interested in taking part with their children contacted the research team directly through a lab email account. Families were invited to participate in the study if parents reported that their child was in the correct age range and had a diagnosis of DCD from a qualified clinical professional and no visual impairment, neurological condition or intellectual disability that would affect this diagnosis. In total, 27 parents arranged a visit to participate in the test battery. Motor difficulties were confirmed in these children by the research team, using the Movement Assessment Battery for Children (MABC-2; [30]). The MABC-2 is a standardised motor assessment battery comprising three components: manual dexterity, aiming and catching, and balance. Children completed a series of age-appropriate timed and non-timed tasks (the 7-10 age band was used for the current study). Raw scores are converted into scaled scores and percentile ranks for each component and for the total score, with total scores below the 5^th^ percentile representing severe motor difficulties, and between 6^th^-15^th^ percentiles representing moderate motor difficulties [30]. Three children scored above the 15^th^ percentile on the standardised test and their data were therefore excluded from the analyses. The impact of the child’s motor difficulties on daily living was also confirmed using the MABC-2 Checklist parent report, with all scores below the 15^th^ percentile. All children completed selected subtests of the Wechsler Intelligence Scales for Children (WISC-IV-UK; [31]) to confirm that no intellectual disability was present. Specifically, one subtest assessing verbal IQ (Vocabulary) and one assessing nonverbal IQ (Matrices) was conducted. Raw scores can be converted into scaled scores (*M*=10, *SD*=3) for each subtest, and any child scoring more than two standard deviations below the mean (i.e. a standard score below 6) on both subtests would be excluded. No children were excluded from the sample based on this criterion. In addition, children were also screened for language impairments using subtests from the Clinical Evaluation of Language Fundamentals (CELF-4-UK; [32]) to potentially divide the sample into groups with or without additional language deficits. Given the small sample size this division was not made, and the data are not reported. The final DCD group therefore included 24 children.

Children were recruited into the TD control group through contacting local schools. A short email was sent to the Headteachers of ten local schools, outlining the study and how the school could get involved. One of these schools responded positively and shared the study information sheets with parents of all children aged 7-10 in the school (*N*=270). Parents who were interested in taking part with their children completed a consent form and returned it to the research team through the class teacher. In total, 36 parents (13%) returned a consent form and their children were tested on the screening measures at school to confirm inclusion in the TD group. One child with a diagnosis of autism chose to participate in the study but was not included in the final analyses. All other children scored above the 15^th^ percentile on the MABC-2 and did not score below a standard score of 6 on both WISC-IV-UK subtests, and no further exclusions were therefore made. Of the 35 remaining children, 23 attended a follow-up visit to complete the experimental task at the University.

Additional background measures assessing participants’ language and attention abilities were also completed by all children completing the experimental task, and any additional diagnoses recorded for those in the DCD group. Inattention and hyperactivity symptoms related to a diagnosis of Attention Deficit Hyperactivity Disorder (ADHD) were assessed through the Conners-3 parent report form [33]. Language abilities were assessed using the Test of Word Reading Efficiency (TOWRE-2; [34]). Total scores are converted to standard scores (*M*=100, *SD*=15). The data from all screening and background measures are presented in Table 1, along with demographic information and additional diagnoses, for participants who completed the full test battery. Children in the DCD group with additional diagnoses were not excluded in order to maximise power and to provide a more representative picture of clinical samples of children with DCD.

**Table 1.**
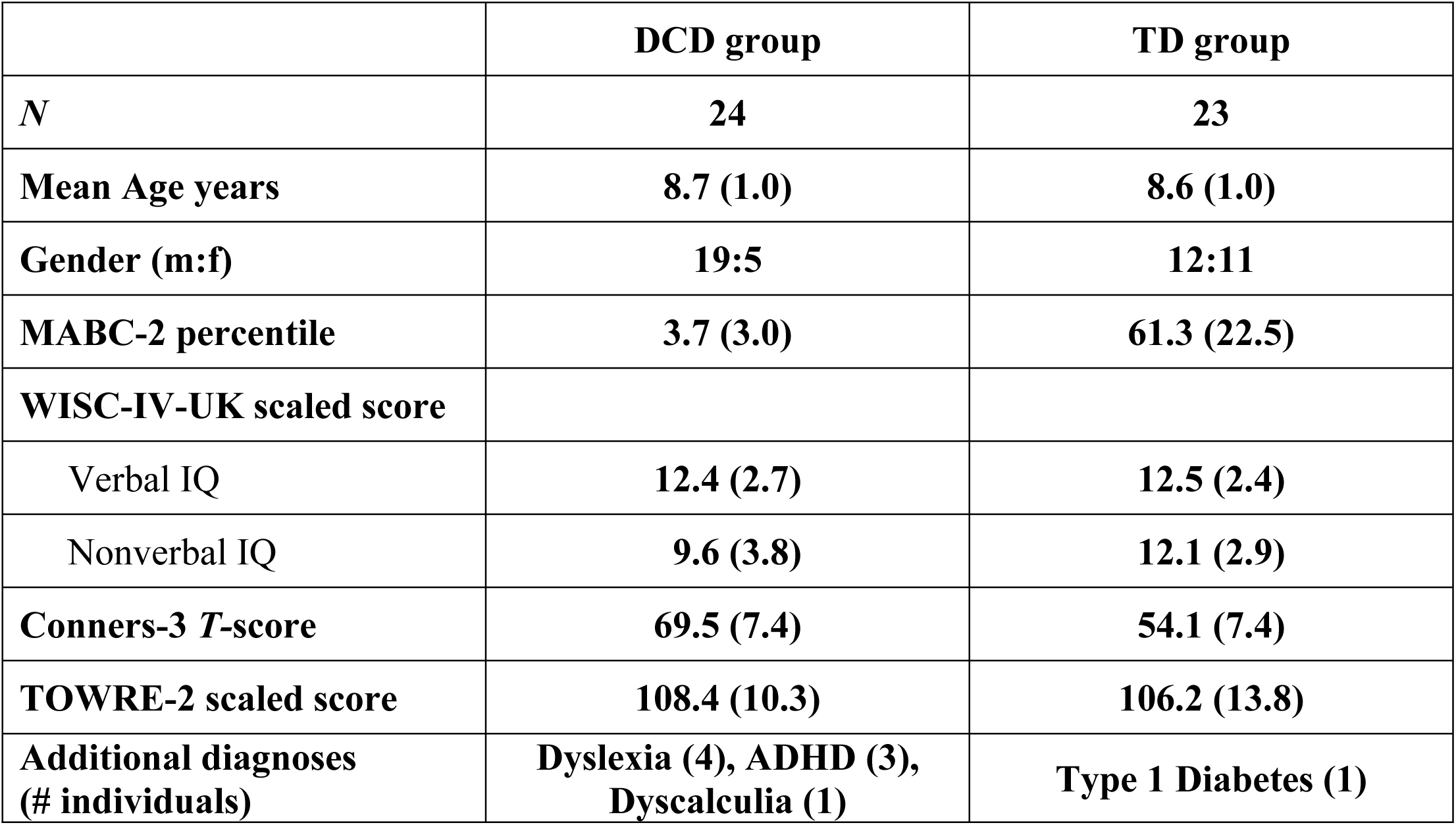
Screening and background information for final DCD and TD groups. Standard deviations in brackets.

### Experimental task

Serial prediction tasks (SPTs) were adapted from Schubotz et al. [18] to produce a child-friendly game in which participants played the role of a ‘sequence detective’. These tasks assessed visuoperceptual, spatial, and temporal processing, with an additional analogous motor task developed for the current study. In the adapted tasks, participants are presented with short sequences of either two objects (visuoperceptual), two object positions (spatial), or two object presentation lengths (temporal) on a computer screen. In the motor task, participants follow sequences by sequentially pressing two areas of a four button-press box each indicated by a flash. In all tasks, one trial consists of six repetitions of these short sequences to enable sequence learning. These short sequences foster the detection of regularities and thus allows to specifically test the internal forward modelling account. In 50% of the trials, the sequence was disrupted towards the end (either 10th or 11^th^ stimulus position within the sequence) by changing the order of stimuli within the sequence (see Figures 1 and 2), while the other half remained unchanged (intact trials). After each trial, participants judged whether the sequence was disrupted at the end by pressing a button on a small separate numerical keypad (identical to the numerical keypad from a standard keyboard) for either ‘yes’ (4) or ‘no’ (6). Behavioural performance was assessed through error rates for this yes/no decision for all tasks. In addition, for the motor task, reaction times (RTs) to produce each element of the sequence were also recorded and analysed. Each task took approximately 10 minutes to complete and task order was counterbalanced, with half of the children completing the tasks in a pre-determined order (Visual, Control, Temporal, Motor, Spatial) and the other half completing them in reverse order.

**Fig 1.**
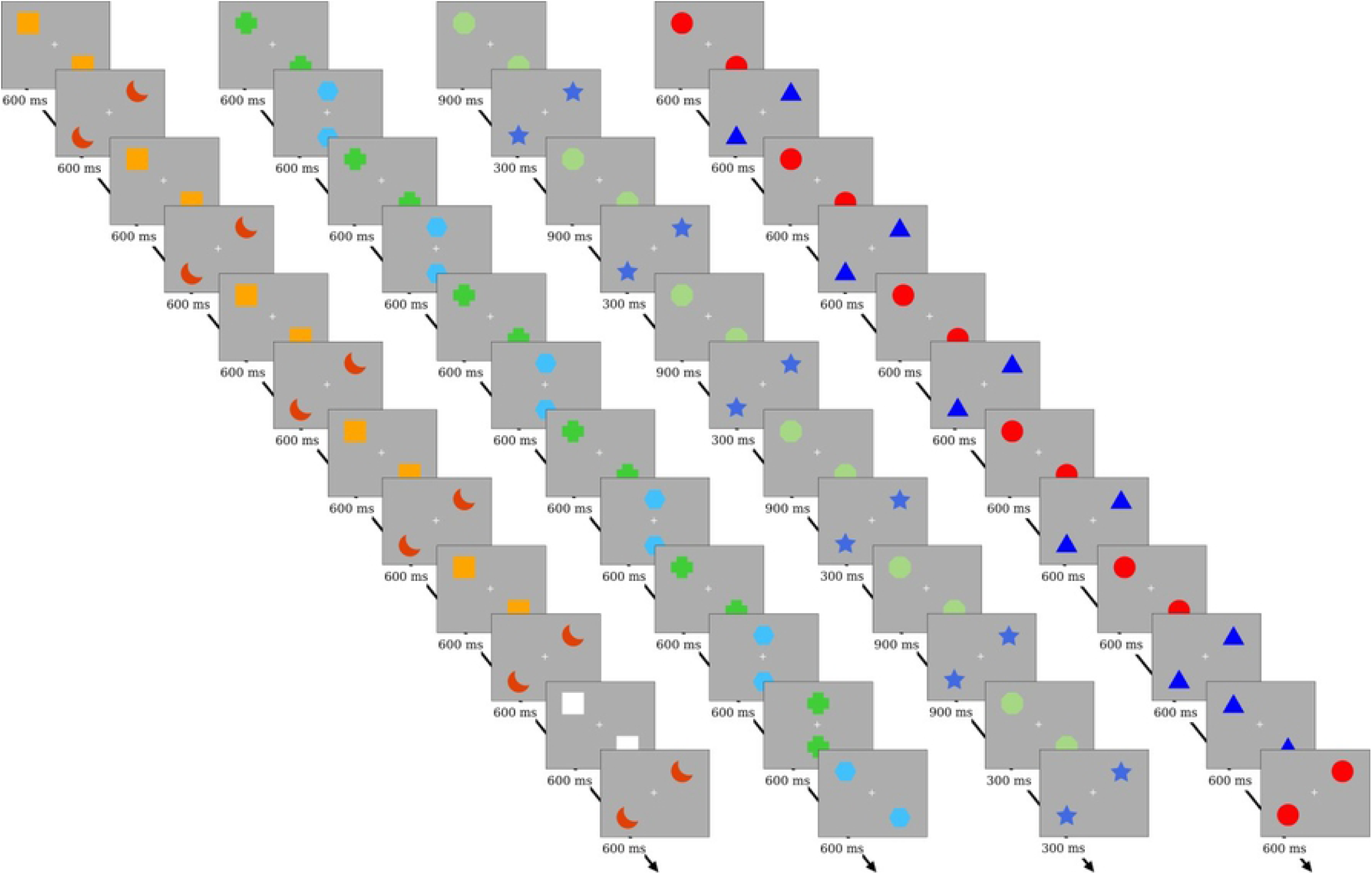
Schematic for a trial in the Sequence Learning Tasks. All show an example of a trial where the sequence is broken. Order of tasks: Control, Spatial, Temporal, Visuoperceptual.

**Fig 2.**
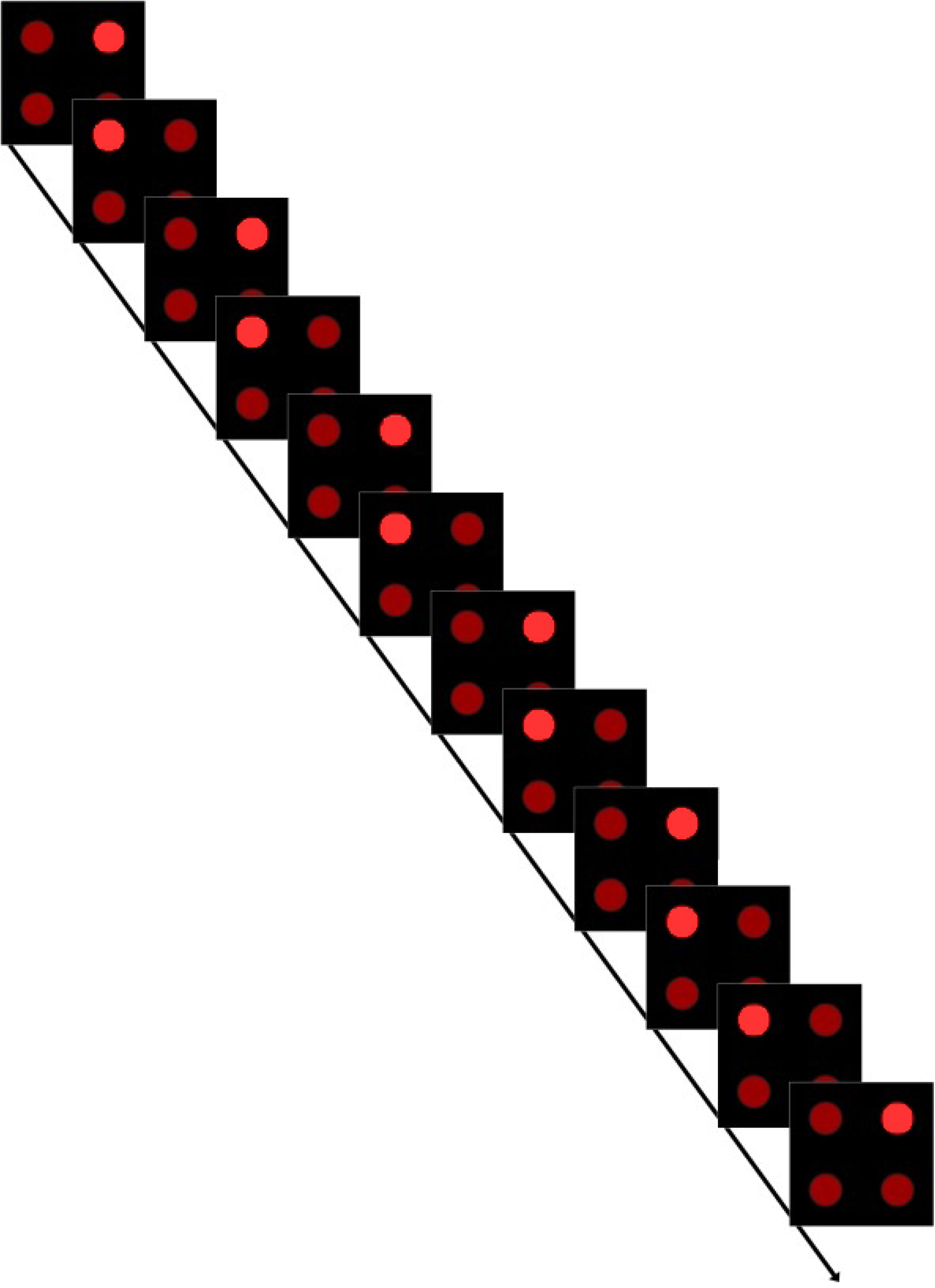
Schematic for a trial in the Motor tasks. In this example the sequence changes. The lights on the response box remained lit until the participant responded.

### EEG recording and preparation

During the experimental tasks, the electroencephalogram (EEG) was recorded using an ActiCap 32 electrode system and recorded using Brain Vision Recorder. The Ag/AgCl electrodes were placed according to the 10-20 positioning standard, with 4 additional external electrodes placed as references (one each on the mastoid bones behind each ear) and to record eye movements (one on the outer canthus of the left eye and the other above the left eye).

### Procedure

Ethical approval was obtained from the University of Surrey Ethics Committee. Children recruited through the local school were visited at school to complete the screening and background measures, with parents asked to complete the Conners-3 questionnaire and send it back anonymously to the research team. Two researchers conducted the test battery, which was split up over several sessions to ensure that children were not absent from class for long periods of time. In the first session, the range of tasks was explained to the children and their assent to participate was obtained. It was explained that more details about each task would be given before they completed it and they would be given a chance to ask questions. They were also told that they could stop at any time and return to their classroom or choose not to complete a task, and that they could stop for breaks whenever they wanted. All children agreed to participate and wrote their name on an assent form to indicate this agreement. All assessments were administered in the same place, a large activity room in the school. The only exception was the throwing and catching subtest of the MABC-2 which was administered in the school playground. Participants completed the MABC-2 in the first session, the WISC in the second session, and the TOWRE in the third session. Approximately a month passed between each session. Children received stickers on the completion of each task.

All data were coded and entered to ensure that children met the inclusion criteria for the TD group, and parents were then invited via email (provided with the initial consent form) to attend the University to complete the experimental task, which was explained in full. All children who completed the background measures received a certificate for their participation, irrespective of whether they took part in the experimental session. Parents accompanied the child to the University for the experimental session after school or at the weekend. On arrival, the procedure was again explained to both the parent and child, including being introduced to the EEG cap and highlighting their right to withdraw, and informed consent was obtained. The EEG cap was then applied. Where the electrode impedance was greater than 10kΩ further preparation was applied (primarily consisting of applying more gel and gently rubbing it in). Throughout this stage the experimenters monitored verbal responses and body language to ensure that the participant was not experiencing undue discomfort with the preparation procedure. Once the EEG application was complete and the child was comfortable, the experimental task was introduced. For each version of the experimental task participants were initially shown an example of a trial where the sequence was not disrupted and at the end of this trial, they were asked to describe the sequence they saw. This was followed by a trial where the sequence was disrupted; the participant was asked if they noticed any difference in that trial. The participants then moved onto practice trials, in which they had to respond correctly three times before they could proceed to the main experimental task. When the participant responded correctly to a practice trial, they received positive feedback, and when they answered incorrectly the trial was repeated at a reduced speed so that they could see where they went wrong. Participants were given short breaks between each task as required, with a longer break (approx. 10 minutes) halfway through where they could move around and have a snack before moving on. The whole battery of experimental tasks took around 90 minutes to complete, including breaks.

For the DCD group, all background and experimental measures were completed during one visit, lasting approximately 6 hours including as many breaks as necessary and lunch. The visit began with obtaining informed consent following the same procedure as for the TD group. Background measures were completed first to allow the child to become familiar with the research team and environment before the experimental procedure. The battery of tasks began with the WISC, followed by the TOWRE, and the MABC-2. This order was chosen to build a rapport between the participant and the researcher and build confidence in the participant before the MABC-2, which was expected to be more difficult for participants in the DCD group. Parents completed the Conners-3 questionnaire and MABC-2 Checklist while their child was completing the tasks. The experimental procedure was then identical to that completed by the TD group. All children in both groups received stickers on completion of each task and a certificate at the end of the visit. During the visit, refreshments were provided, and travel expenses were reimbursed. No other incentives or compensations were provided.

## Data analyses

### Behavioural data

#### Sequence prediction/reception

Raw data were processed in R Studio (Version 1.2.1335; R version: 3.6.3) in order to extract accuracy scores for each sequence type (disrupted vs. intact) on a participant-by -task basis. These scores were used to calculate each individual’s *d*-prime score for each task. Participants with fewer than 10 valid trials for either of the conditions were excluded from the analyses. These data were then submitted to a multivariate analysis of variance. The *d*-prime score for each of the conditions was used as the dependent variable and Group membership was used as the independent variable.

#### EEG data

Raw EEG data were processed in Matlab (Version 2017a) using the EEGLAB toolbox (Version 14.1.2). The data were split into the sections corresponding to each task for pre-processing. The data were filtered with a Hamming windowed sinc FIR filter with a band-pass set at 0.1 Hz to 40 Hz. The filtered data then underwent independent component analysis (ICA) in order to identify and manually remove blinks from the EEG data. A window of -250 to 1000 milliseconds relative to a stimulus onset marker was used to epoch the data. For trials in disrupted sequences, the stimulus selected for epoching was the first stimulus where the sequence was disrupted. For intact trials, the data were epoched around the stimuli in the same positions (i.e. either stimulus 10 or 11 in the sequence). A baseline of -250 ms to 0 ms was used for all trials.

After the data were epoched, trials with artefacts or noise in the data were removed using a semi-automated system. First, trials where any of the channels passed beyond a ±100uV threshold were automatically identified. This was followed by manual inspection and rejection of trials. Finally, any participants with fewer than 10 trials in either of the conditions were excluded from the analysis.

Individual trials were averaged by sequence type to produce the ERP for each sequence type for each participant and condition. These ERPs were then analysed using non-parametric permutation statistics, with cluster correction for multiple comparisons (Fieldtrip [35]). This analysis would reveal clusters of adjacent time points for which the waveforms to disrupted and intact sequences differ significantly from each other.

Differences in the neural responses to the sequences (disrupted vs. intact) were first investigated within each group separately by comparing the waveforms averaged across CP1 and CP2 electrodes for these two sequence types. In order to investigate whether there were any differences in the responses between groups, a difference wave was produced for each group by subtracting waveforms to disrupted and intact trials. These difference waves were compared to one another to ascertain whether the difference in neural responses differed by group (the interaction between group and sequence type).

## Results

### Demographic and Background Variables

The two groups were first compared on demographic and background variables (cf. Table. 1). Both groups were of the same age, *t*(45) < 1, *p* = .73 but had a different gender distribution, χ^2^=7.0, *p*=.012. In contrast to the TD group, there were significantly more boys than girls within the DCD group. A similar gender ratio of children diagnosed with DCD in the general population has previously been observed [36]. Children in the DCD group had poorer motor abilities as indicated by a significant lower percentile on the MABC-2, *t*(45) = 13.7, *p* < .0005 and higher inattention and hyperactivity symptoms as indicated by higher scores on the Conners-3 questionnaire, *t*(45)=6.9, *p*<.0005. They had also lower non-verbal IQ, *t*(45) = 2.9, *p* = .01 but showed similar verbal IQ, *t*(45)<1, *p*=.88 compared to the TD group. Both groups were also comparable regarding their word reading efficiency, *t*(45) < 1, *p* = .54.

### Sequence prediction across domains

#### Behavioural accuracy

As is apparent from Fig. 3, the DCD group showed lower *d*-prime scores than the TD group in all conditions aside from the control. The MANOVA with factors Group and condition revealed a significant main effect of Group, *F*(5, 21) = 7.35, *p*= .0004. Follow up ANOVAs revealed significant main effects of group for all conditions aside from the control condition: *F*(1,25) = 1.03, *p*= .32 (control task); *F*(1,25) = 4.52, *p*= .04 (motor task); *F*(1,25) = 9.51, *p*= .005 (spatial task); *F*(1,25) = 8.95, *p*= .006 (temporal task); *F*(1,25) = 23.30, *p*= .00006 (visual task).

**Fig 3.**
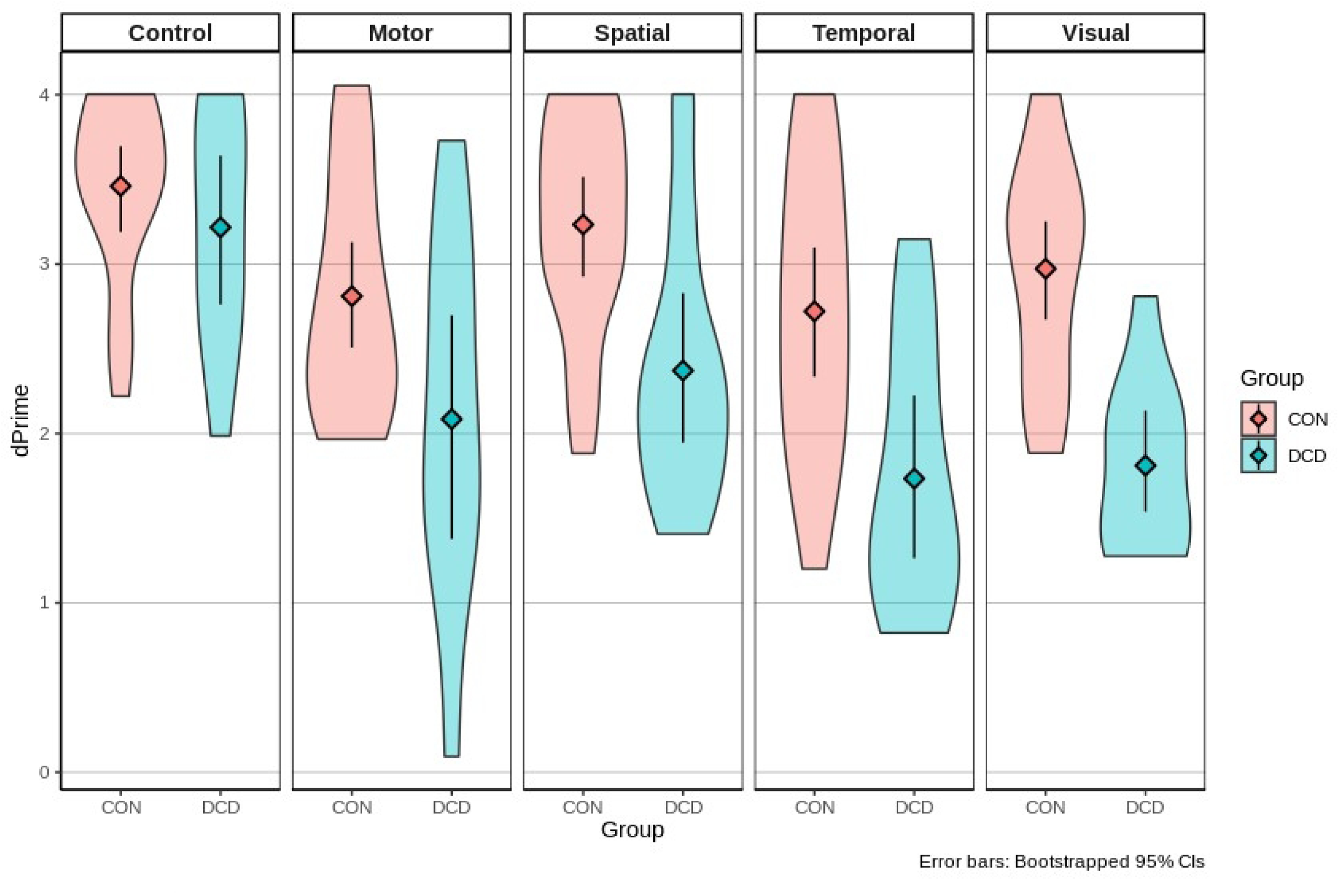
Behavioural accuracy for trials with and without sequence disruptions for children with Developmental Coordination Disorder (DCD) and typically-developing (TD) controls in perceptual and motor serial prediction tasks.

In the three perceptual SPT tasks, the same analyses revealed significant main effects of group, *F*(1,42)=29.75, *p*<.001, η_g_^2^=0.27 [visual]; *F*(1,39)=26.07, *p*<.001, η_g_^2^=0.25 [spatial]; *F*(1,41)=21.99, *p*<.001, η_g_^2^=0.24 [temporal], with the DCD group being less accurate than the TD group across tasks. In addition, both groups had lower accuracy on the disrupted compared to intact sequences, with main effects of sequence across tasks, *F*(1,42)=17.43, *p*<.001, η_g_^2^=0.17 [visual]; *F*(1,39)=11.41, *p*=.002, η_g_^2^=0.13 [spatial]; *F*(1,41)=42.87, *p*<.001, η_g_^2^=0.30 [temporal]. Finally, and in contrast to the control task, significant group x sequence interactions in all tasks revealed that the DCD group compared to TD group performed more poorly when sequences were disrupted than when they were not, *F*(1,42)=4.86, *p*=.03, η_g_^2^=0.05 [visual]; *F*(1,39)=6.60, *p*=0.01, η_g_^2^=0.08 [spatial]; *F*(1,41)=5.03, *p*=.03, η_g_^2^=0.05 [temporal].

The same analyses conducted on accuracy in the visuomotor task also revealed a significant main effect of group, *F*(1,39)=6.27, *p*=.02, η_g_^2^=0.08, with lower accuracy in the DCD group than the TD controls. However, there was no significant main effect of sequence, *F*(1,39)=3.36, *p*=.07, η_g_^2^=0.04, and no significant interaction between variables, *F*(1,39)=0.14, *p*=.71, η_g_^2^=0.002.

#### Neural responses to sequence disruption

Both groups exhibited more positive-going waveforms for stimuli disrupting the sequence compared to their respective counterparts in intact sequences in all conditions. This difference between disrupted and intact sequences seem to emerge at around 400ms post-stimulus onset over central parietal regions, resembling a P3 component of the ERP (see Fig. 4).

**Fig 4.**
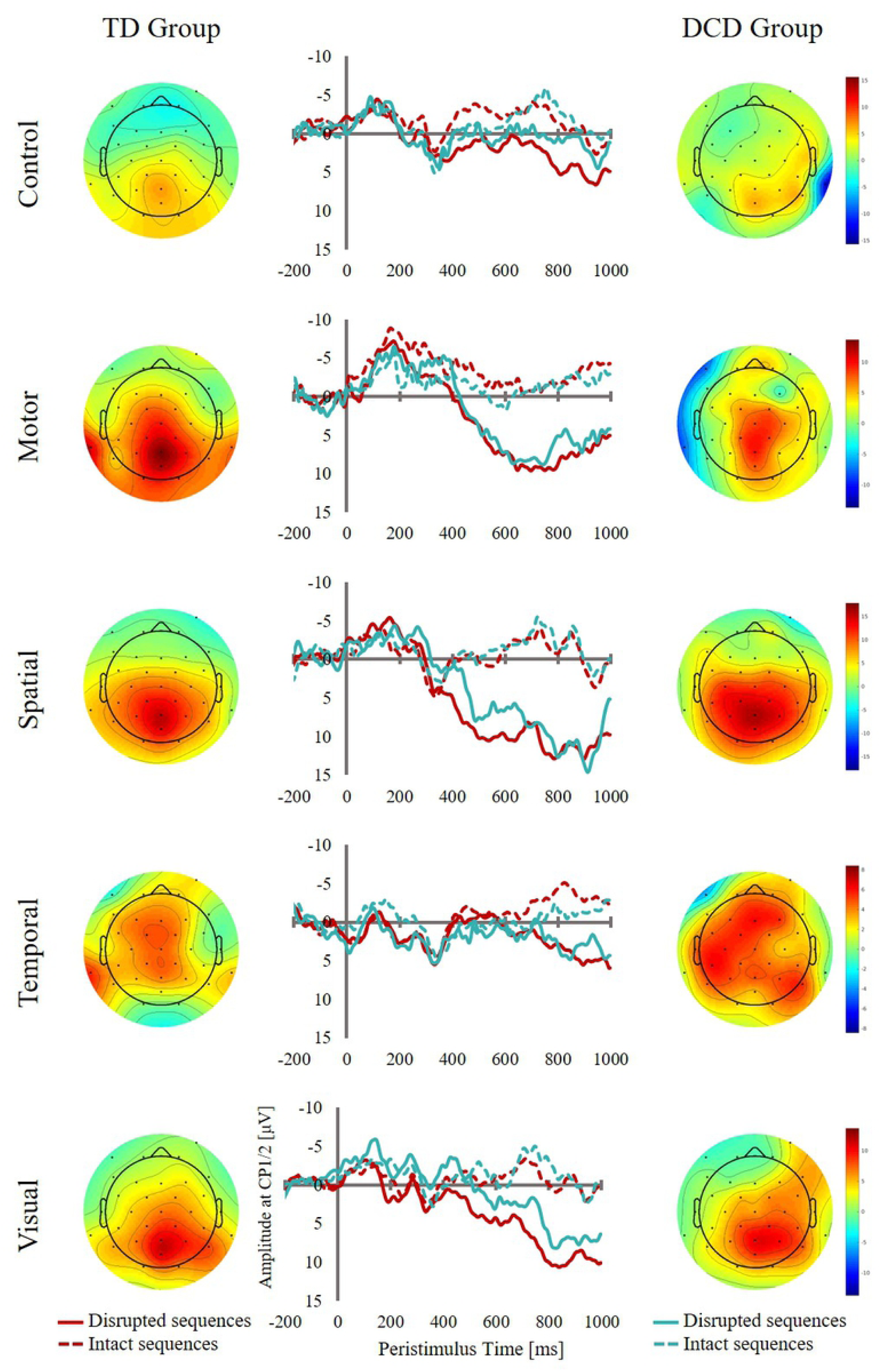
ERP waveforms collapsed across CP1 and CP2 electrodes (middle) and topographical maps for DCD (right) and TD groups (left). The maps show activity averaged for the 500 – 900ms and the 700 – 900ms time windows for TD and DCD groups, respectively.

Non-parametric permutation statistics were computed separately for each condition to reveal temporal clusters where this difference reached significance for the two groups. In the control task, different response patterns for disrupted and intact sequences were observed when the groups were analysed separately. For the TD group the observed data showed a significant effect of sequence type in the control task for two clusters; the first cluster extended from 376ms to 620ms (p = 0.008) and the second extended from 630ms to 998ms (p = 0.001). In both clusters the disrupted sequence showed more positive amplitudes than the intact sequence. In contrast, for the DCD group no statistically significant clusters were observed between the two conditions.

In the motor task again two clusters showed a significantly more positive waveform for disrupted compared to intact sequences for the TD group; the first cluster extended from 496ms to 694ms (p = 0.007) and the second extended from 774ms to 998ms (p = 0.003). The main effect of sequence type for the DCD group did reach significance in a cluster extending from 716ms to 998ms (p = 0.005). Similar to the control task, the interaction between group and sequence type did not reach significance.

In the spatial task a similar picture emerged. For the TD group an extensive cluster spanning the entire time window from 370ms to 998ms showed a significant effect of sequence type (p = 0.0007), with disrupted sequences exhibiting a more positive-going waveform compared to intact sequences. For the DCD group only one cluster from 504ms to 932ms showed the same significant effect of sequence type (p = 0.004). The interaction between group and sequence type did not reach significance.

The same pattern of results was obtained for the visual task. Again, the TD group showed more positive-going waveforms for disrupted sequences in an extensive time window ranging from 450ms to 1000ms (p = 0.0007). For the DCD group, a later cluster was observed extending only from 750ms to 950ms. The permutation test revealed that, within that cluster, there was a statistically significant effect of sequence type (p = 0.009). As for all other tasks, the disrupted sequence showed more positive amplitudes than intact sequences.

For the temporal task the TD group showed a significant effect of sequence type only in the later time window extending from 700ms to 998ms (p = 0.0007), while the DCD group showed no such effect.

Taken together, the TD group showed one extended cluster of significant differences in amplitude elicited by disrupted and intact sequences across all tasks starting around 500 ms across all tasks. This difference was not significant for a brief time window in the control and motor tasks only. Based on timing and scalp topography, this difference between disrupted and intact sequences resembles a P3 component. In contrast, the DCD group showed significant differences between disrupted and intact sequences only between 750ms and 900ms for three out of five tasks: the visual, spatial and motor tasks.

#### Learning in the visuomotor task

In order to assess the learning of the motor sequence in the visuomotor task, the reaction times for responses for each trial were extracted and analysed (see Fig. 5). Group differences between the median reaction times for the very first responses were compared using a *t*-test in order to assess whether there were any differences in overall response speed. The *t*-test indicated that there was not a significant group difference, *t*(37.47) = -1.51, *p*= .14). Additionally, the reaction times to stimuli disrupting the sequence were compared to their respective counterparts in intact sequences. An ANOVA with the factors sequence type (disrupted vs. intact) and group (DCD vs. TD) revealed a main effect of sequence type (*F*(1,43)=107.53, *p*<.0005, η_g_^2^=0.312) but no main effect of group (*F*(1,43)=1.85, *p*=.181, η_g_^2^=0.034) or an interaction between the two factors (*F*(1,43)=0.87, *p*=.357, η_g_^2^=0.004). This indicates that reactions times were longer for both groups for disrupted compared to intact sequences.

**Fig 5.**
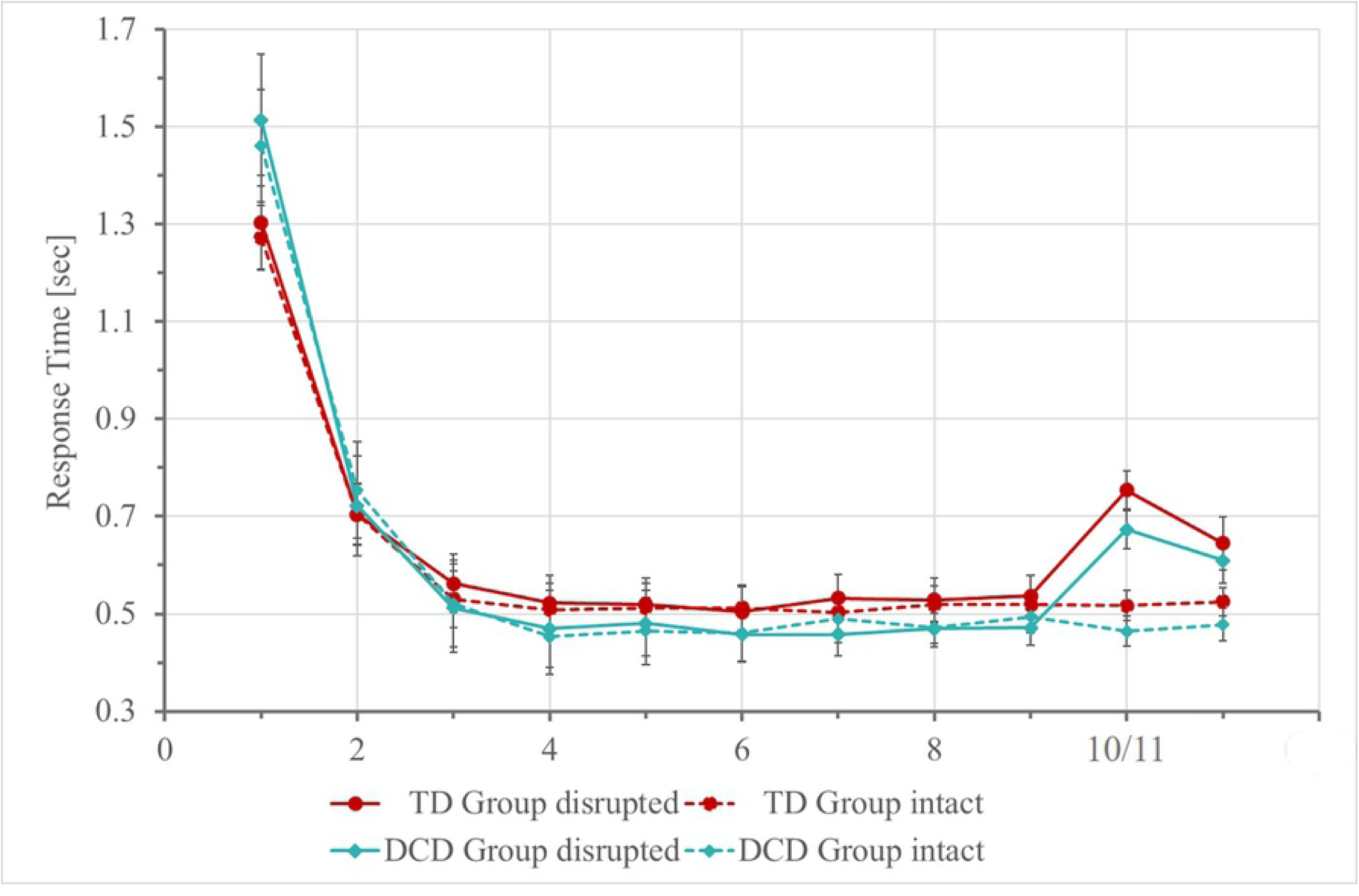
Reaction times in the motor sequence learning condition for children in the DCD (blue) and TD (red) groups. Data points indicate the mean of median reaction times for each stimulus in disrupted (solid lines) and intact (dotted lines) sequences. Error bars represent standard errors. NB: Stimuli 10/11 indicate reaction times for the stimuli disrupting the sequence and their respective counterparts in intact sequences.

In order to assess whether there were any group differences in the slopes of the learning curves, a linear model was fitted to the log-transformed reaction times, with response number as the predictor variable. The estimated slopes were then submitted to a *t*-test, which revealed a significant difference between the slopes, *t*(42.5) = 2.61, *p*= 0.01. The DCD group had a more rapidly-decreasing slope than the control group (mean slope in DCD = -0.098, mean slope in Control = -0.075).

Taken together, these results do not provide any evidence that children with DCD are impaired on implicit motor sequence learning. Specifically, they demonstrated a similar improvement in reaction times when learning the motor sequence, as well as a similar slowing for trials disrupting the sequence, as typically-developing children.

## Discussion

The current study aimed to better understand sequence learning in visuomotor tasks in DCD, for which previous research has provided mixed results. Based on the internal forward modelling account of DCD, the study specifically investigated sequence prediction abilities in children with DCD across motor and non-motor domains. Using serial prediction tasks (SPTs), both sequence perception and sequence production were assessed to further disentangle which stage of the internal forward model is affected in DCD. The study focused specifically on children to reflect the previous procedural learning research in DCD. Investigating these issues as early as possible in development, in the time that DCD is usually diagnosed, is important for timely intervention.

Behavioural results indicated that children with DCD improved their reaction times during motor sequence learning but showed poorer explicit discrimination of intact and disrupted sequences. Crucially, this impaired explicit discrimination was observed across different domains, including the visuospatial and the temporal domain. This was accompanied by the absence or a severe delay of the P3 component of the event-related potential in all tasks for children with DCD. Crucially, this delay occurred during the implicit processing of sequence disruptions and before the explicit decision on whether the sequence was in fact disrupted was made. In accordance with previous research implicating this ERP component in stimulus evaluation processes and the updating of a mental model of the environment [29], this suggests a domain-general impairment of sequence prediction rather than a selective motor impairment.

### Implications for an internal forward modelling account of DCD

The present results seem to be in line with the forward modelling account of DCD [15,16]. They suggest that across several perceptual domains children with DCD are unable to evaluate incoming stimuli with sufficient accuracy in real time. This deficit in stimulus evaluation and updating implies that, for sequences with individual elements that occur in fast succession, the evaluation of an element is not yet completed when the next element needs to be processed. This could result in a ‘blurred’ representation of the sequence elements hindering the extraction of regularities and, consequently, the build-up of an accurate internal forward model and the precise prediction of the next element. Crucially, the deficit observed in the present study affects both motor and non-motor domains, thereby confirming previous studies demonstrating deficits in children with DCD across several experimental measures assessing planning, inhibition, working memory and cognitive flexibility (see [37] for an overview). In the motor domain, this inaccurate internal forward model would cause feedback from the environment to be inefficient so that high levels of accuracy can only be obtained at the expense of very slow and careful responding. This view is in agreement with previous findings demonstrating reduced general processing speed in children with DCD [38,39]. The study by Bernardi and colleagues [39] reported that children with DCD were most impaired in a visual scanning task requiring them to find and cross out all examples of the number ‘3’ presented amongst a visual array of different numbers and letters. This could be interpreted in light of the present findings as reflecting impaired stimulus evaluation and inefficient sequencing of the visual scanning. In non-motor domains, such deficits in updating the mental model would become evident in all tasks requiring such updating, as in working memory or inhibitory control tasks. For instance, during response inhibition participants are required to update a prepotent response with an alternative response. A delay in updating, as suggested by the present results, would then necessarily result in the slower response times found in several previous studies on response inhibition in DCD [e.g., 12,13]. The slowing of response times reported in working memory tasks in DCD [37,38,41] could also be explained by this delay in updating: these tasks typically entail the constant updating of the content held in memory in response to dynamic stimulus presentation.

### Implicit vs explicit sequence learning in the visuomotor task

Although the SRTT is often considered to demonstrate implicit learning by improved RTs to the sequences over the course of the task, this conclusion has been debated (e.g., [42]). In particular, the task has been criticised because of the difficulty in assessing whether the participants are explicitly aware of the sequence they are learning, and at what stage they become conscious of this fact. Previous research has yielded mixed results [22,23]. Both studies used a sequence generation test to assess explicit awareness of the sequence structure. While Gheysen and colleagues [22] reported sequence generation to be at chance level, Lejeune et al. [23] reported at least partial explicit awareness. Furthermore, it is possible to be aware that a sequence is present but to have difficulty reproducing the full sequence after the task; thus, the distinction between implicit and explicit learning in the SRTT is not as clear as suggested. The use of the serial prediction task (SPT) in the present study allows us to further delineate explicit sequence learning. Our results suggest that children with DCD are able to explicitly discriminate between intact and disrupted sequences across all domains tested, albeit their discrimination is significantly lower than for typically-developing children. This might suggest that the previous results reported from the SRTT are mainly driven by a diminished ability in DCD to detect and learn the statistical regularities in sequential material, rather than a pure motor planning/execution deficit. The apparent heterogeneity of results in previous studies could also be reconciled by considering differences in response mode required between the standard SRTT and the current implementation of the SPT paradigm. There are two apparent differences to take into account: (i) the current custom-made response box requires far less precision of the movement of one or both hands (according to participants’ preference), while in the standard SRTT bimanual responses of index and middle fingers are required, and (ii) the assignment of the screen position to a corresponding key has to be kept in working memory for the SRTT but not for our version of the SPT (see [23] for a similar argument). Thus, the response mode of the standard SRTT requires sequencing of the motor response and thus puts high demands on updating the mental forward model. By reducing updating demands for the motor response during task performance in the present study, the results seem to reveal an intact ability for implicit motor learning.

## Conclusions

To conclude, the present results suggest an impairment in DCD of the updating of an internal forward model. This may result in a blurred representation of that model and, consequently, in a reduced ability to detect regularities in the environment (e.g., sequences). Although the behavioural results could also be explained by an impaired motor realisation of the predicted sequence, electrophysiological brain responses (i.e., the delay in the P3 component) support the above interpretation of deficits in the updating of an internal forward model. Crucially, this deficit affects motor and non-motor domains to a similar degree. A further implication of the present study, in line with previous suggestions [23], is that implicit motor adaptation during sequence learning can be observed in DCD when the updating demands for the required motor response are low. It should be noted, however, that the present sequence is rather short, therefore future research in this area should continue to explore more complex sequences. In these complex sequences the detection of regularities might be more difficult and, thus, other mechanisms might be more important for sequence learning. Further developing our knowledge of internal forward modelling in DCD could be vital for our understanding of the wide range of co-occurring difficulties experienced by those with a diagnosis.

